# Nutrients and flow shape the cyclic dominance games between *Escherichia coli* strains

**DOI:** 10.1101/2022.08.15.504033

**Authors:** Thierry Kuhn, Junier Pilar, Redouan Bshary, Céline Terrettaz, Diego Gonzalez, Xiang-Yi Li Richter

**Affiliations:** Institute of Biology, University of Neuchâtel, Rue Emile-Argand 11, 2000 Neuchâtel, Switzerland

**Keywords:** 3D printing, Bacterial community dynamics, Bioreactor, Evolutionary game theory, Individual-based simulation, Rock-paper-scissors game

## Abstract

Evolutionary game theory has provided various models to explain the coexistence of competing strategies, one of which is the rock-paper-scissors (RPS) game. A system of three *Escherichia coli* strains—a toxin-producer, a resistant, and a sensitive—has become a classic experimental model for studying RPS games. Previous experimental and theoretical studies, however, often ignored the influence of ecological factors such as nutrients and toxin dynamics on the evolutionary game dynamics. In this work, we combine experiments and modeling to study how these factors affect competition dynamics. Using 3D-printed mini-bioreactors, we tracked the frequency of the three strains in different culturing media and under different flow regimes. Although our experimental system fulfilled the requirements of cyclic dominance, we did not observe clear cycles or long-term coexistence between strains. We found that both nutrients and flow rates strongly impacted population dynamics. In our simulations, we explicitly modeled the release, removal and diffusion of toxin. We showed that the amount of toxin that is retained in the system is a simple indicator that can predict competition outcomes across broad parameter space. Moreover, our simulation results suggest that high rates of toxin diffusion might have prevented cyclic patterns from emerging in our experimental system.

## Introduction

Cyclic dominance interactions represent an interesting type of evolutionary dynamics that have been suggested to promote species coexistence in nature [1–4]. In contrast to hierarchical competition dynamics, where stronger competitors tend to exclude weaker ones, in cyclic dominance interactions, there is no obvious best competitor. Each player can beat some competitors, but loses when confronted to others, like in the children’s “rock-paper-scissors” game, where paper wraps rock, rock crushes scissors, and scissors cut paper, forming a dominance circle (Figure 1A). This type of interaction seems rare in animals (yet with famous examples, such as the mating strategy polymorphism in side-blotched lizards [5]), but is more common in plants [6–8] and widespread in the microbial world. Cyclic dominance might evolve more easily in microorganisms than in multicellular eukaryotes because of their large population size and high per capita mutation rate [9]. Moreover, in bacteria, competition is frequently driven by bactericidal toxins whose distribution and target ranges are highly variable; cyclic dominance will tend to arise whenever competing strains are sensitive to the toxins produced by some but not all other competing strains [1,10–12].

**Figure 1.**
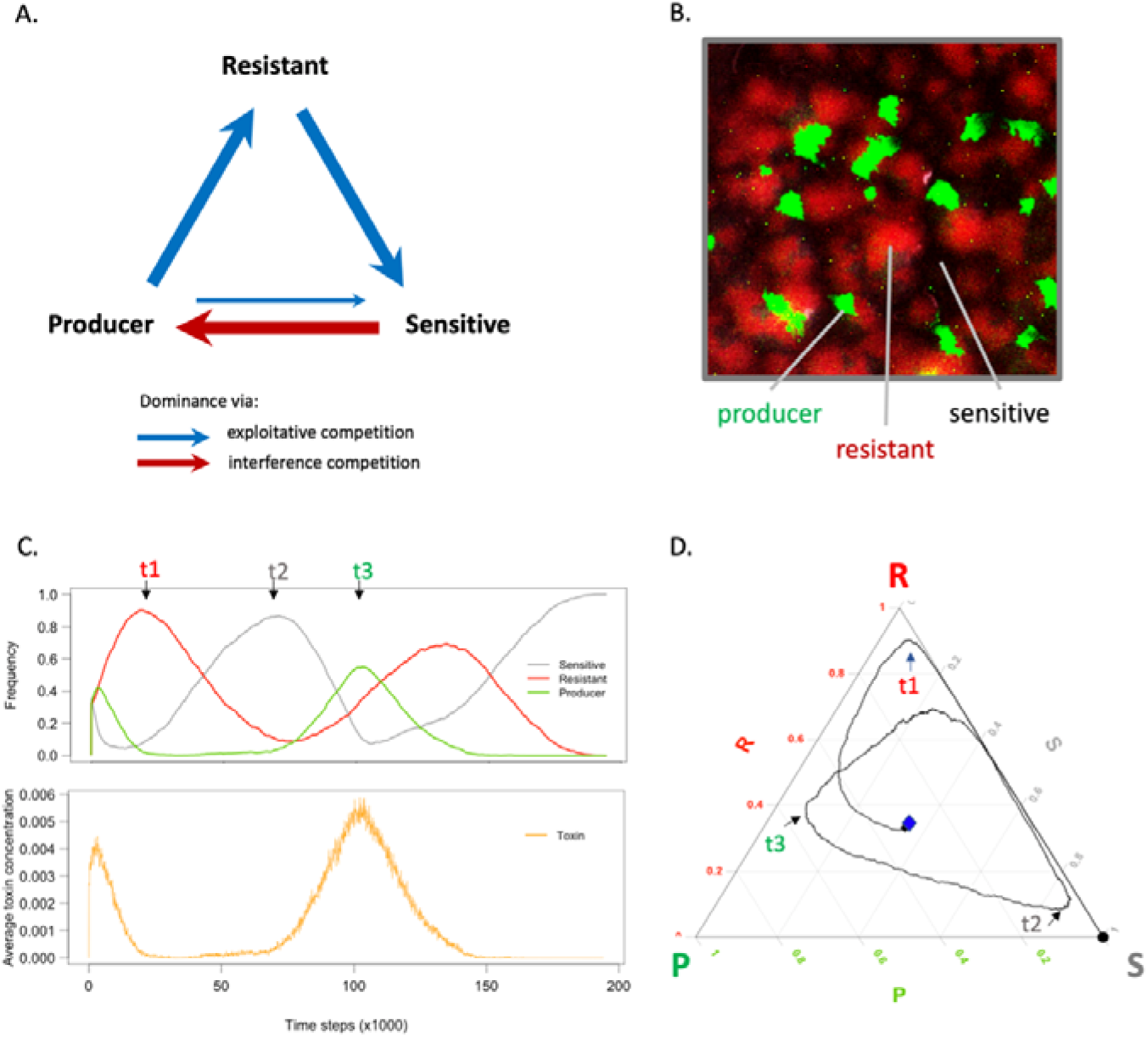
Basic concepts underlying the research. A. Cyclic dominance driven by toxin-mediated killing (i.e., interference competition [23,24], represented by the red arrow) or higher growth rate (i.e., exploitative competition [23,24], represented by the blue arrows) between three *E. coli* strains. B. Micrograph of a multi-strain biofilm on a nutrient agar plate. The producer, resistant and sensitive strains are in green (GFP), red (DsRedExpress) and black, respectively. C. Changes in strain frequency (upper part) and toxin concentration (lower part) as a function of time during a simulation run of the model. In the upper graph, green, red, and grey lines represent the frequencies of the producer, the resistant, and the sensitive strains over time; in the lower graph, the orange line represents the toxin concentration (which correlates, with some delay, with the frequency of the producer strain). D. Alternative representation of the strain frequencies shown in panel C as a “ternary plot.” The strain frequency changes are represented as a trajectory between three vertices representing each one of the three genotypes: P for the producer, R for the resistant, and S for the sensitive strain. The three timepoints t1, t2, and t3 match between panels C and D.

*Escherichia coli* is a mammalian gut commensal and pathogen that heavily relies on toxins, collectively named colicins, to exclude conspecific competitors. This bacterium has been used as a model system for studying cyclic dominance both experimentally and theoretically. In the gut, strains that produce colicins, strains that are sensitive to colicins, and strains that, at some cost, have acquired resistance to colicins often coexist [13,14]. In such a system, the toxin producers kill sensitive individuals, the resistant individuals outgrow the toxin producers, and the sensitive individuals outgrow the resistant ones in pairwise competitions, which suggests that a “rock-paper-scissors” evolutionary game might underlie the coexistence between the strains (Figure 1AB). Mathematical models and experiments based on this system have identified important factors that determine the likelihood of strain coexistence, including the range of spatial interaction [2], mobility rate [4], spatial heterogeneity [15], and community size [16]. However, previous models and experiments have often neglected the role of some key ecological factors at play in *E. coli’s* natural environment, which can dramatically change the competition dynamics between the players in the “rock-paper-scissors” evolutionary game.

The mammalian gut is a complex and dynamic ecosystem in which parameters like pH, nutrient availability, spatial structure, and flow constantly vary, both locally and globally [17]. Some of these parameters, especially nutrients and flow, may influence the competitive interactions between *E. coli* strains by affecting their relative growth rates and the local concentrations of colicin. Because colicins are protein-based toxins, their production is particularly costly and inefficient in environments where amino acids are limiting [18]. The production of colicins is therefore often directly modulated by nutrients [19–21]. The concentration of toxins is also highly sensitive to hydrodynamic factors. Due to their high molecular weight (40 to 80 kDa [18]), the spatial and temporal distributions of colicins can be affected by their varying diffusion rates in the heterogeneous intestinal extracellular matrix [22]. Moreover, the rate of toxin removal, by flow or degradation, can change throughout the gut. Because colicins are potent and fast-acting [18], small changes in toxin concentration could dramatically impact the sensitive strain survival and threaten coexistence between the three strains. However, the effects of toxin removal have rarely been studied as such in mathematical models and are usually not controlled or accounted for in experiments [2,12].

In this work, we combined laboratory experiments and computer simulations to study how nutrients and toxin production and removal rates affect the “rock-paper-scissors” evolutionary game in *E. coli*. Our experiments involved continuous cultures of communities of the three *E. coli* strains in 3D-printed mini-bioreactors with different media compositions and flow rates. The bioreactor we designed captures some important features of the mammalian gut by allowing bacterial cells to form biofilms on the bottom of the culturing chamber, resembling the microbial community on the gut epithelium. In addition, we can control the content and speed of medium flow into the bioreactor, which allows us to simulate different gastrointestinal conditions. To help discriminate between toxin-related and growth-related effects, we built a simulation model that independently controls toxin production, toxin diffusion, and toxin removal on the one hand, and growth parameters on the other hand. Through stepwise simulations, the model follows population dynamics and toxin levels in a spatially tractable way (Figure 1CD; see also Figure 3AB). By combining experiments and simulations to study cyclic dominance in a model system of three *E. coli* strains, we gained knowledge on how nutrients and flow shape the competition outcomes in a highly controlled system, which may further help us understand how biodiversity in microbial communities is maintained in natural environments.

## Materials and Methods

### 1. The experiments

We use three strains of the bacterium *E. coli:* the BZB1011 pColA strain (labeled with a green fluorescent protein) that produces colicin A; the BZB1011 *btuB* strain (labeled with a red fluorescent protein) that does not produce the toxin but is resistant to it; and the BZB1011 strain (unlabeled) that neither produces the toxin nor is resistant to it [25]. The bacterial strains were inoculated with a 1:1:1 ratio and cultured at 37°C for 14 days in mini-bioreactors (Figure 2) of 1 ml working volume that we designed and 3D-printed. The medium was constantly renewed through an inflow connector, connected to a medium dispenser, and an outflow connector, connected to a waste container; the flow of medium was controlled by two pumps, one for the inlet, and one for the outlet. Sterile air supply was ensured by an aeration pipe connected to a 0.22 μm mesh filter. A funnel-shaped sampling chamber capped by a rubber stopper served for inoculation and sampling purposes. We implemented a factorial design of three different culturing media with high, intermediate, and low casamino acid to glucose (CA:G) ratios and two flow rates for a total of six different experimental conditions, each with five independent replicates. We followed the community dynamics in each bioreactor by taking a sample each day and quantifying the relative frequencies of the three strains. In addition, we measured the growth rates of the three strains in monoculture and the toxin production rates of the producer strain in all three media. We also experimentally confirmed that the producer strain can rapidly kill the sensitive strain thanks to secreted colicins.

**Figure 2.**
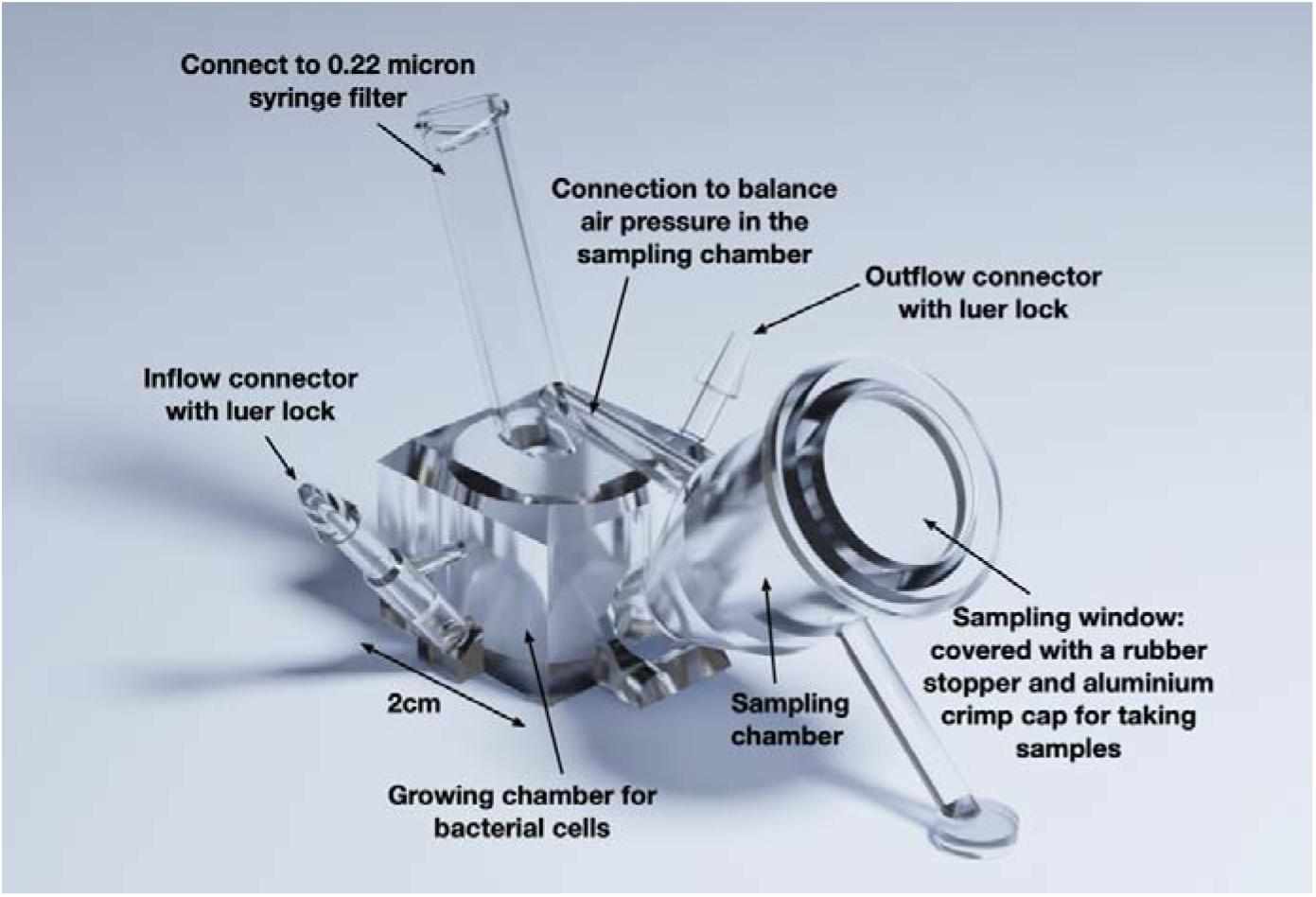
A 3D illustration of the mini-bioreactor used for continuous cultures of the bacterial communities in the experiments.

Detailed CAD drawings of the mini-bioreactor design and the associated STL file for 3D printing are provided in the ESM. We also provide detailed information on the experimental procedures and quantification methods in the Supplementary Materials and Methods.

### 2. The simulation model

To help better understand the experimental results, we designed a spatially explicit, individual-based simulation model. Our model was based on a lattice with periodic boundaries, where each position can either be empty or host a bacterial cell (i.e., the May-Leonard formulation [26,27]) and contains information about the local toxin concentration (Figure 3). We considered three bacterial cell types, corresponding to the colicin-producing, resistant, and sensitive strains of *E. coli* we used in the experiments. To initiate the simulation, we populated 1% of the positions in the lattice with a bacterium chosen at random among the three strains. The toxin concentration was set to zero everywhere. In each of the subsequent time points, we updated every position of the lattice following the order of (i) cell death and removal, (ii) toxin concentration dynamics (including the production, diffusion, and removal of colicin), and (iii) colonization of the empty positions by cell growth. The simulations were stopped either when coexistence was lost (i.e., only one cell type was left) or when they reached 300’000 steps. A detailed description of the simulation model is provided in the Supplementary Materials and Methods, and the annotated simulation codes are available in the ESM. The model parameters needed for understanding the *Results* are listed in Table 1.

**Figure 3.**
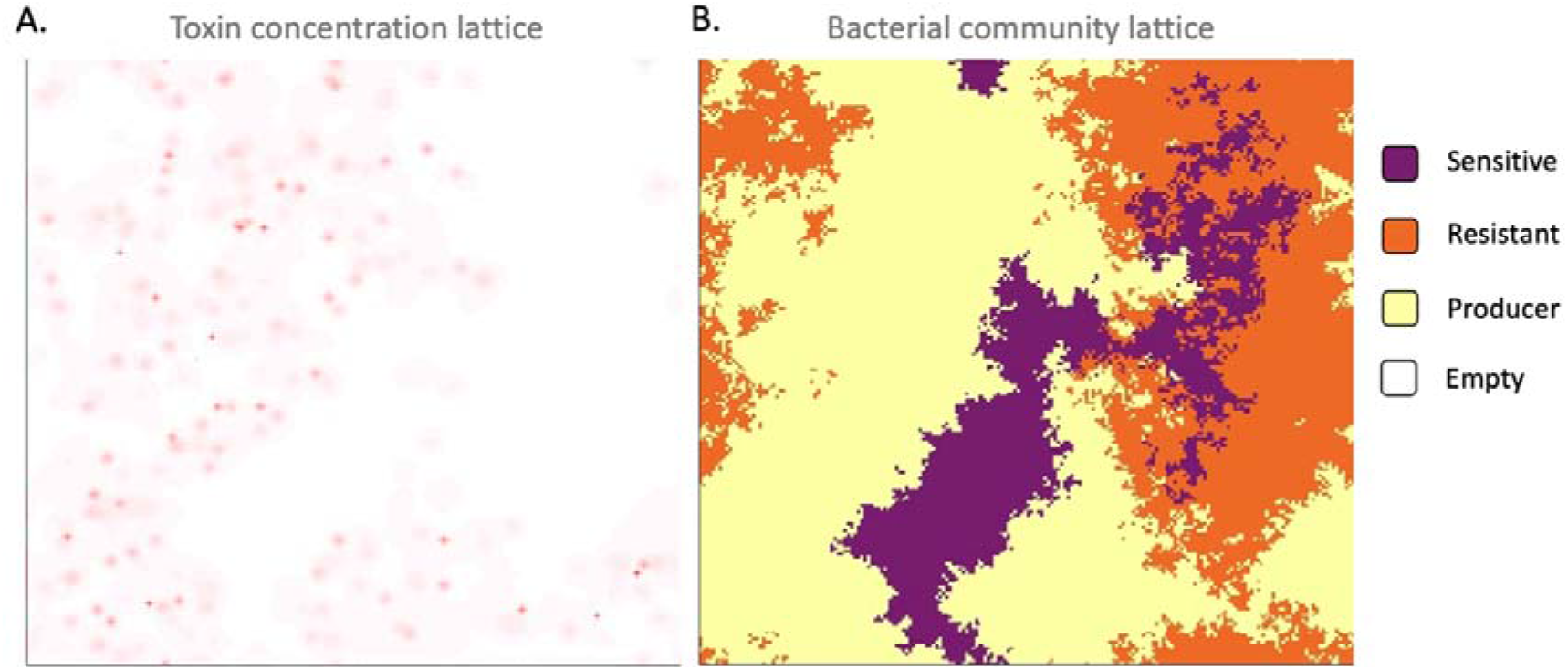
An illustration of the simulation model. Panels A and B show the toxin concentration and spatial occupation of bacterial cells in the lattice, respectively. The state of the simulation corresponds to timepoint t3 in Figure 1CD.

**Table 1.** Parameters implemented in the simulation model that are mentioned in the *Results*.

|  |  |
| --- | --- |
| $D$ | Diffusion rate of colicin |
| $u$ | Removal rate of colicin by the flow of culturing medium |
| $d_l$ | Spontaneous lysis probability of the toxin-producing cells per time step |
| $p$ | Amount of toxin released when a producer cell lyses |
| $v_R$ | Cost on growth rate for colicin resistance |
| $v_P$ | Cost on growth rate for colicin production |

## Results

### 1. Experimental results

#### 1.1 Behavior of the three E. coli strains fitted a “rock-paper-scissors” game

Before studying the competition dynamics between the three *E. coli* strains, we first confirmed that their behavior in monoculture fitted the requirements of a “rock-paper-scissors” game in the three different media used (“high CA:G,” “intermediate CA:G,” and “low CA:G”). In particular, we evaluated the relative growth rate of the three strains and the toxin production of the producer strain in the three media. We found that in all nutrient conditions, the sensitive strain had the highest growth rate, followed by the resistant strain, and by the colicin-producing strain. All pairwise comparisons between the growth rate distributions were still significant (*p* < 0.001) after using the Bonferroni correction for multiple comparisons (Figure 4A). The average cell doubling times of the *E. coli* strains are shown in Supplementary Figure S1. Because of the relatively low nutrient availability in all the culturing media, the doubling times were longer than the typical 20 minutes in rich medium but probably shorter than in the wild [28]. While strains grew faster with higher CA:G ratio, the relative growth rate difference between the toxin-producing and sensitive strains decreased as the CA:G ratio increased in the medium.

**Figure 4.**
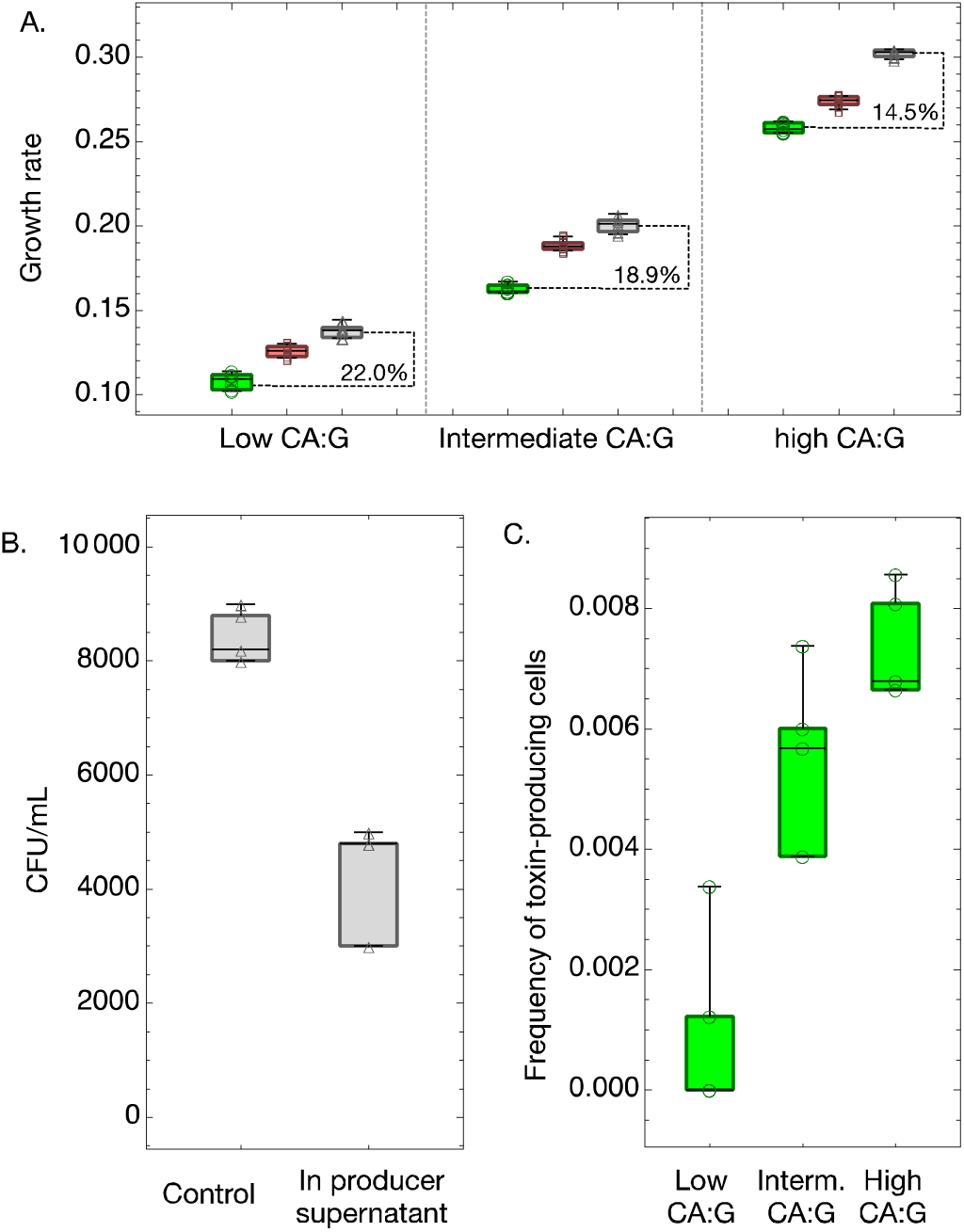
The formal conditions for cyclic dominance are fulfilled in our experimental system. A. The growth rates of three *E. coli* strains under different nutrient conditions. The box-whisker charts filled with green, red, and gray colors and marked with circle, square, and triangle markers represent the toxin-producer, the resistant, and the sensitive strains, respectively. B. Addition of 50% of filtrated supernatant from the producer strain reduces the number of viable sensitive cells by more than 40% in 30 minutes. The panel shows the number of surviving sensitive cells after exposure to sensitive (control) or producer supernatant. C. The frequency of toxin producing cells in a toxin-producer population under different nutrient conditions. The boxplot center lines show the median, box limits show upper and lower quartiles, whiskers show the 1.5x interquartile range, and each marker corresponds to the result of an independent replicate. We have eight, four and four replicates for each box-whisker chart in panels A, B and C, respectively.

We confirmed that the producer strain could kill the sensitive strain via a secreted toxin by exposing sensitive cells to the supernatant of producer cells for 30 minutes (Figure 4B). The relative toxin production of the producer strain in the different media was evaluated using a colicin A transcriptional reporter. We could observe that the frequency of producer cells actively releasing colicin was about 1.5-fold and 10-fold higher in the “high CA:G” medium compared to the “intermediate CA:G” medium and the “low CA:G” medium respectively (Figure 4C). Overall, this indicates that, in our system, nutrients have a major impact on the toxin production rate and the toxin concentration. Based on the relative growth rates of the three strains and the toxin production of the producer strain, the conditions for cyclic dominance as shown in Figure 1A are expected to be fulfilled in all three media. However, because toxin production correlates with the proportion of CA in the medium, the advantage of the producer strain over the sensitive strain is expected to be much stronger in “high CA:G” than in “low CA:G”.

#### 1.2 Effects of nutrient and flow regime on the competition dynamics of E. coli strains

We next followed the population dynamics of the three *E. coli* strains in the three different media and under two different flow regimes (slow and fast) for two weeks in the 3D-printed mini-bioreactors. We found consistent effects of the nutrient condition and the flow regime on the community composition trajectories (Figure 5). In the high CA:G ratio medium, the producer dominated the competition, making up more than 80% of the bacterial population over the entire duration of the experiments after the first day, but neither the resistant nor the sensitive strain became extinct. In contrast, the resistant and sensitive strains appeared more competitive in the medium with low CA:G ratio, where the producer never reached more than 10% of the population. In the medium with intermediate CA:G ratio, the competition outcomes were variable among the replicates, suggesting that the overall competitivities of the three strains were closer in this nutrient environment.

**Figure 5.**
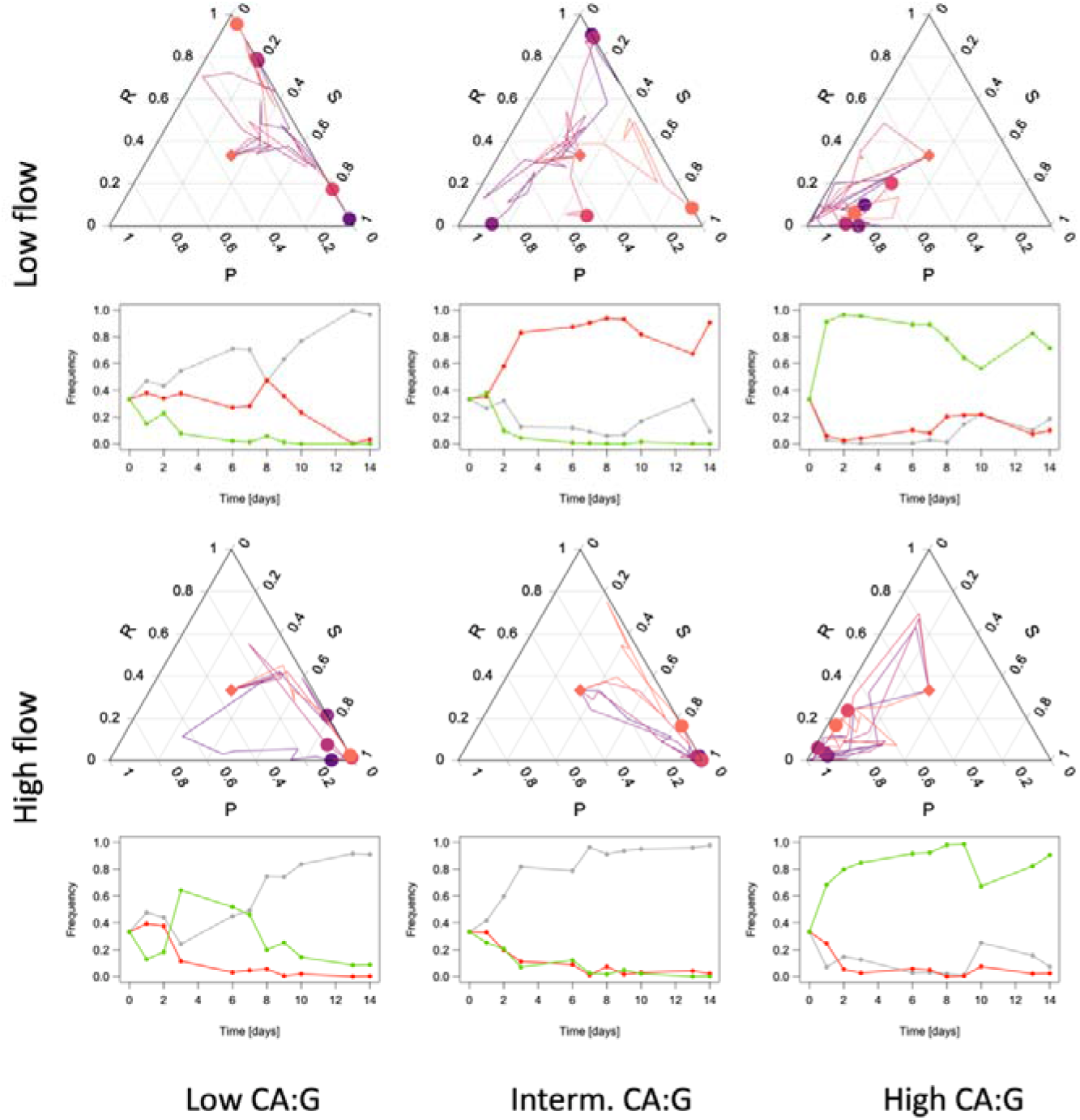
Ternary plots of the community composition of three *E. coli* strains over two weeks in three different media and under two different flow regimes. The frequency of each strain at each time point can be read on the respective side of the ternary plot. Five independent trajectories were plotted in different colors under each condition. The starting point of all trajectories is marked by a diamond; the end point of each trajectory is marked with a circle. Under each ternary plot, one of the five trajectories (the one in the darkest purple hue) is plotted on a linear scale; the sensitive, resistant, and producer strains are shown in grey, red, and green respectively.

Besides the marked effect of the nutrient environment, the effect of the medium flow rate on the competition dynamics was also noteworthy. In the medium with low CA:G ratio, the flow rate determined whether the resistant or sensitive strain dominated the competition in the time window of our experiments: while a higher flow rate favored the sensitive strain, a lower flow rate favored the resistant strain. A higher flow rate could indeed give an advantage to the sensitive cells by removing the toxin more quickly. The competition dynamics under low flow rate in the medium with intermediate CA:G ratio were most interesting: all replicates followed a consistent community composition trajectory over time, but the trajectories of different replicates strongly diverged from one another. This suggests that the slightly different initial community composition caused by stochasticity in the inoculation or during the first hours of the experiments may have strongly impacted the subsequent evolutionary trajectories. This could happen if the overall competitiveness of the three strains were in a much closer range under this experimental condition than in the other conditions we tested.

### 2. Simulation results

#### 2.1 Slower toxin diffusion tends to promote strain coexistence, but up to a limit

To explore in more detail the impact of diffusion, flow, toxin production, and growth rate on the population dynamics of the three *E. coli* strains, we designed and implemented a spatially explicit model of bacterial toxin-based cyclic competition. We first assessed the impact of toxin diffusion on the coexistence of the three competing strains and on the competitivity of each strain. The simulations tended to produce longer coexistence times when slower toxin diffusion rates were used (Figure 6; Supplementary Figure S2). Very short coexistence and no cyclic dominance were characteristic of the instantaneous toxin diffusion case, which is equivalent, when it comes to the toxin concentration, to a well-mixed environment (Figure 6, *D* = 1.0). Because, in this case, sensitive cells are killed at random over the entire lattice, the toxin gives the same advantage to the resistant as to the producer cells; since the resistant cells grow faster than the producer, the resistant wins whenever the toxin reaches concentrations sufficient to eradicate the sensitive and the sensitive wins whenever they do not. With slower diffusion rates, the sensitive population becomes less affected overall by the same amount of released toxin because it takes more time for it to reach patches of sensitive cells; however, in this case, the dying sensitive cells are more often located at the interface with producer patches, which helps the latter strain conquer new space and gives it a colonization advantage. Nevertheless, below a certain toxin diffusion threshold, coexistence breaks down again: the toxin does not diffuse fast enough to free space around the growing producer patch and to help its population increase in frequency. Coexistence is therefore only possible at intermediate toxin diffusion rates. Besides coexistence, diffusion also impacts the tentative competition outcomes: while high diffusion rates (*D* = 1.0) favor the resistant strain, low diffusion rates (*D* = 0.0001) favor the sensitive; intermediate-low (*D* = 0.002) diffusion rates increase the likelihood of the producer winning the competition.

**Figure 6.**
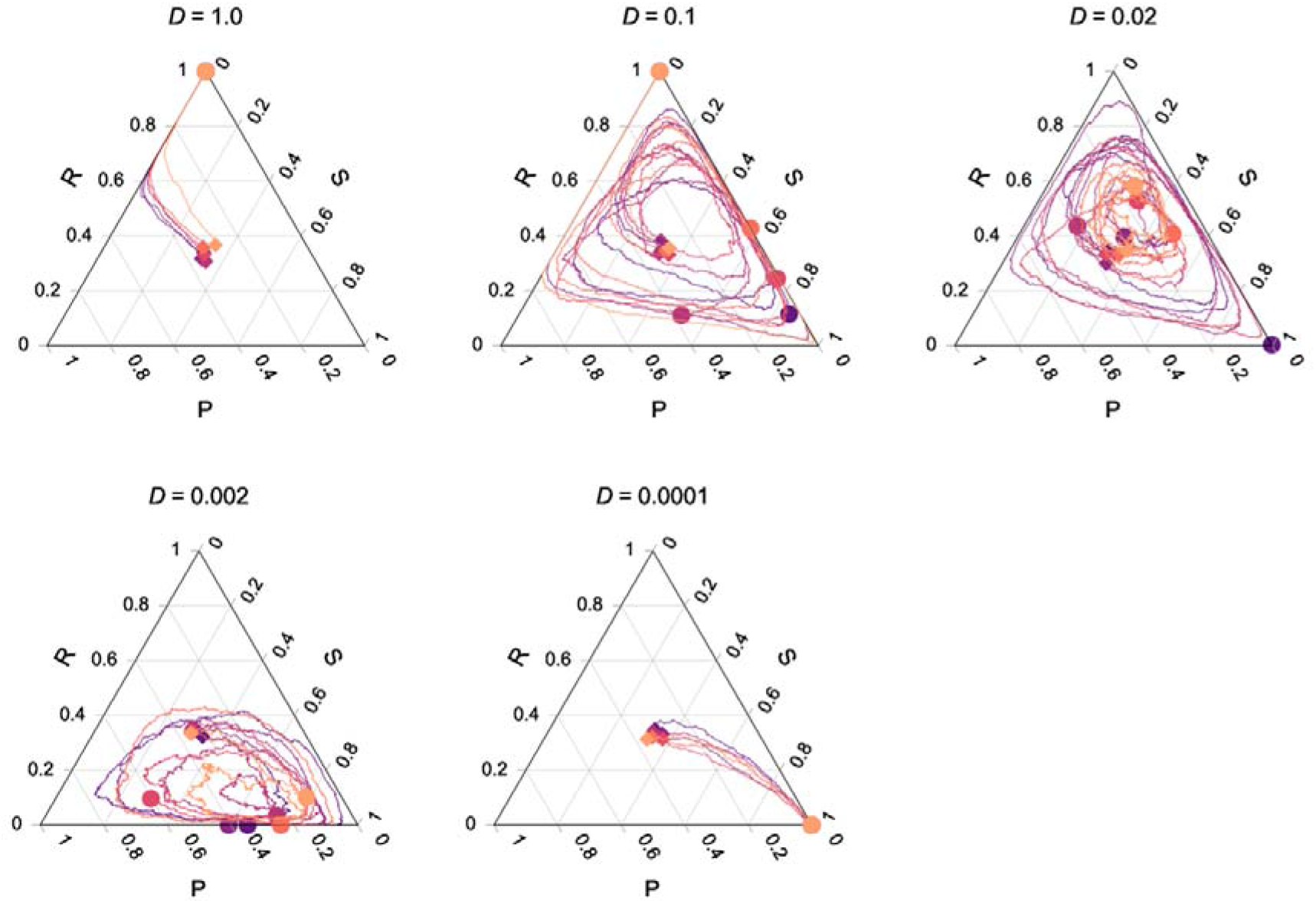
Competition dynamics between the three *E. coli* strains at different toxin diffusion rates (*D*). Very high (*D* = 1.0) and very low (*D* = 0.0001) toxin diffusion rates were detrimental to the colicin producing strain. At intermediate diffusion rates, the frequencies of the three *E. coli* strains fluctuate in cycles that can be more or less favorable to one of the three strains. The spontaneous lysis rate of the colicin producers and the removal rate of colicin were set to intermediate values: *d_l_* = 10^-4^, and *u* = 0.015. Six independent simulation trajectories were plotted in different colors under each condition. The start point and end point of each trajectory are marked with a diamond and a circle of the same color as the trajectory, respectively.

#### 2.2 Strain coexistence is favored at intermediate toxin accumulation

The levels of toxin in the system are controlled by the producer lysis rate (*d_l_*), which correlates with toxin release, and by the toxin removal rate (*u*) through medium flow. Both parameters can impact the concentration of toxin in the system, as can be observed in a community consisting of the colicin producer strain only (Figure 7A; Supplementary Figure S3): decreasing *d_l_* and increasing *u* both lead to lower toxin levels in the system. Although toxin does not reach a steady state in populations comprising all three strains because the producer frequency fluctuates over time, the amount of the toxin contributed by each producer cell which is retained in the system (or “toxin retention”), *pd_l_* (1 – *u*), hereafter denoted as *c*\*, nonetheless controls toxin accumulation in the system. Our simulations showed that long-term coexistence is favored at intermediate *c*\* levels. When the values of *c*\* were too high, like in the cases of low or medium flow rates (*u* = 0.0075 or *u* = 0.015) in combination with high toxin release rates (*d_l_*=3e-4) (Figure 7B), most of the sensitive cells were killed at the very beginning of the simulations runs, and the producer was subsequently outcompeted by the resistant strain (or by the sensitive one if some cells happened to survive the initial colicin wave). In contrast, when the values of *c*\* were too low, like in the cases of high or medium flow rates (*u* = 0.015 or *u* = 0.03) in combination with low toxin release rate (*d_l_* = 3e-5) (Figure 7B), the colicin producers were not able to maintain a toxin level that was high enough to kill sensitive cells. Consequently, the sensitive strain, which has the highest growth rate, excluded the other two strains. Only when *c*\* was kept within an intermediate range, cyclic population dynamics could emerge, and the coexistence of the strains could be maintained for a relatively long duration.

**Figure 7.**
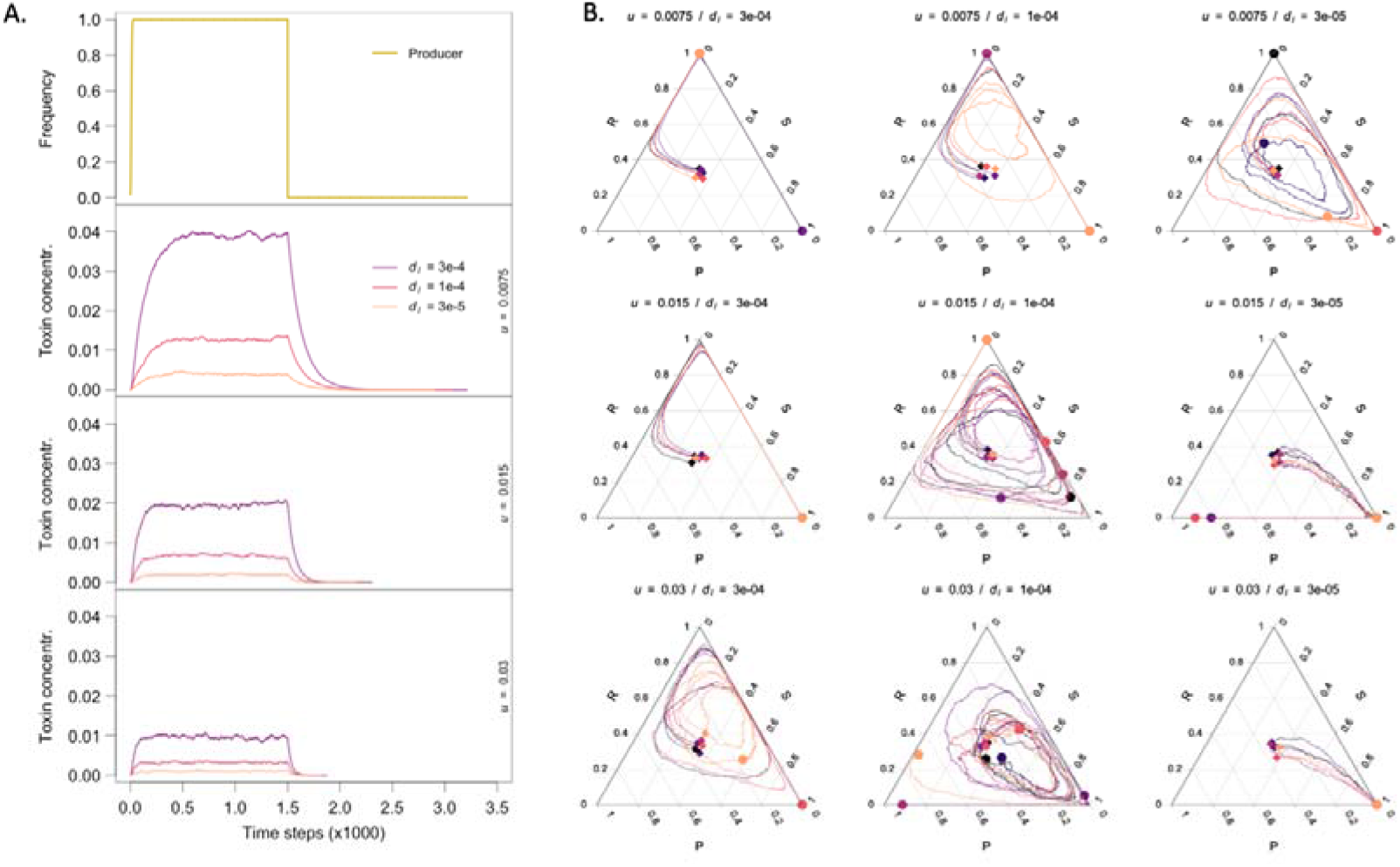
The impact of toxin accumulation on population dynamics. A. Toxin accumulation and toxin removal in a producer population at different toxin production and removal rates. A producer population was left to occupy the entire space for 1500 time steps and then eliminated; toxin increase and decrease (until it reaches the 1e-7 threshold) is observed. Three elements emerge that could affect the strain population dynamics: (1) the toxin steady-state level (plateau), which is the resultant of both production and removal rates and seems to have the highest impact on coexistence, (2) the toxin accumulation rate when the producer population is growing, mainly impacted by the toxin production rate, and (3) the toxin concentration decrease rate when the producer population plummets, mainly impacted by the toxin removal rate. B. Population trajectory for the nine combinations of toxin production and removal rates we used in panel A. The toxin diffusion rate is set to fast (*D* = 0.1); the plots for instantaneous (*D* = 1.0) and slow (*D* = 0.02) diffusion rates can be found in Supplementary Figure S4 for comparison. There are six independent simulation trajectories under each condition, indicated by lines of different colors. The start point and end point of each trajectory are marked with a diamond and a circle of the same color as the trajectory, respectively. Toxin production rate *d_l_* = 3e-5, 1e-4, 3e-4; toxin removal rate *u* = 0.0075, 0.015, 0.03.

#### 2.3 Extreme toxin fluxes can lead to population fluctuations that break down coexistence

Although the lysis rate of the colicin producers *d_l_* and the removal rate of toxin *u* can both influence the accumulation of colicin in the system, their effects on the competition dynamics between the bacterial strains are not entirely interchangeable. The former controls how fast colicin can accumulate in the system (Figure 7A, colicin concentration rose faster at higher *d_l_* values); the latter controls how fast colicin can be flushed out (Figure 7A, colicin concentration returned to zero faster at higher *u* values). At the bacterial population level, *d_l_* determines the speed of the sensitive strain collapse when the producer frequency starts to increase, whereas *u* determines if and when the sensitive strain can bounce back from a toxin wave after the producer’s frequency has declined. To show the effects of *d_l_* and *u* without confounding effects, we adjusted these two parameters correlatively while maintaining the toxin retention, *c*\*, constant (Figure 8; Supplementary Figure S5). At too low and too high *d_l_* and *u* values, long-term coexistence between the strains was compromised. In the former case, the toxin was not removed fast enough at the onset of the competition and the sensitive strain was heavily hit by the initial toxin wave leading to the dominance of the resistant strain (or rarely, after a bounce-back, the sensitive strain) over the producer. At too high *d_l_* and *u* values, the toxin could not be accumulated to a high enough concentration to threaten the sensitive population at the onset of the competition, which then favored the sensitive strain (or sometimes the producer) at the expense of the resistant strain. Such extreme toxin fluxes, through their effect on the sensitive population, increase the probability for one of the three strains to be outcompeted at the very beginning of the competition and for coexistence to break down. Under intermediate toxin production and removal rates, by contrast, the fluctuations of toxin levels were damped, which limited the risk of global disbalance that it entails. Both toxin accumulation and toxin flux need therefore to be within an intermediate range to support long-term strain coexistence.

**Figure 8.**
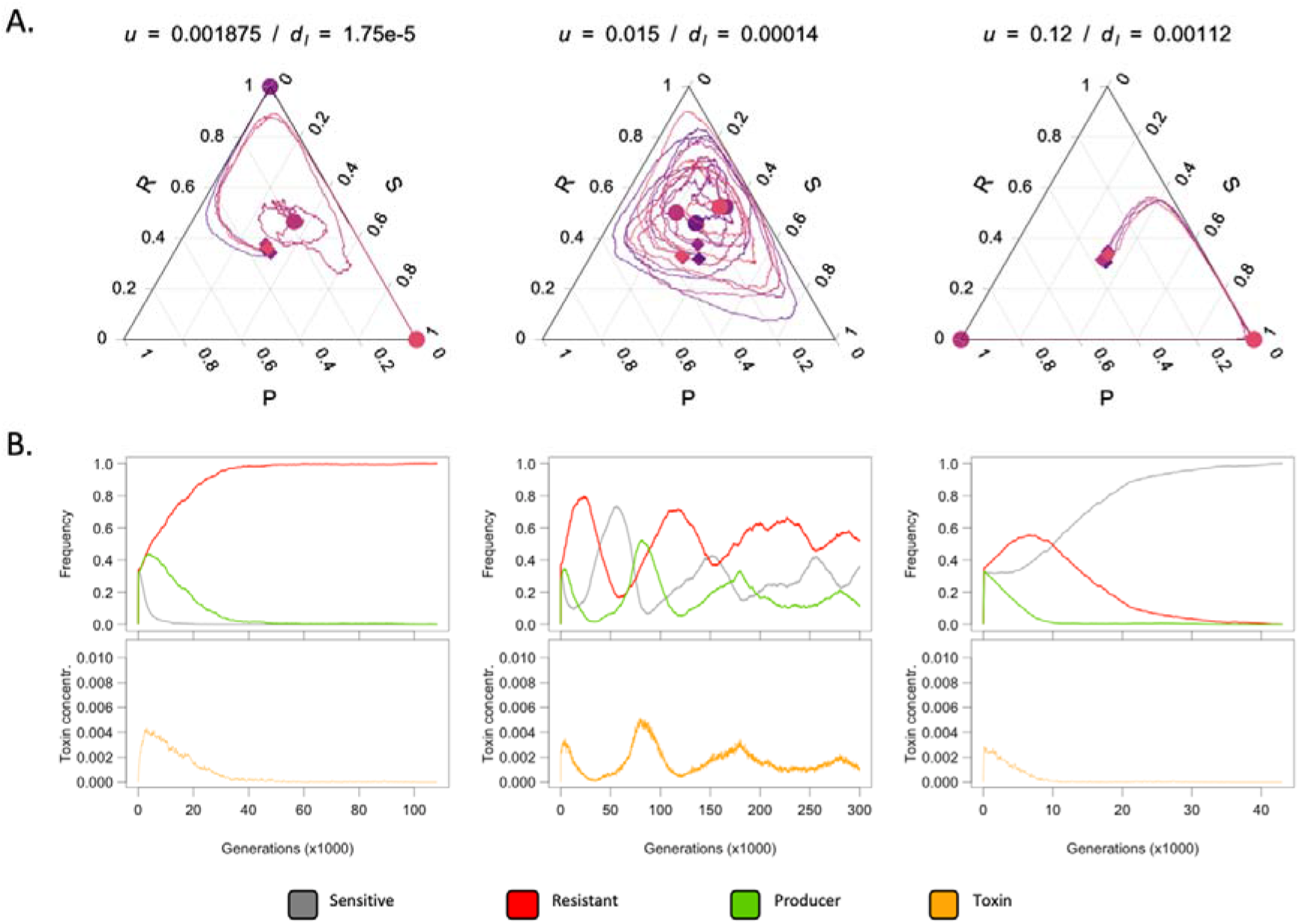
The impact of toxin flux on population dynamics. A. Impact of increasing flow and toxin production on cyclic competition patterns with the toxin retention, *c*\*, kept constant. There are four independent simulation trajectories under each condition, indicated by lines of different colors. The start point and end point of each trajectory are marked with a diamond and a circle of the same color as the trajectory, respectively. B. Dynamics over time of the three strains and the toxin levels for one of the four replicates. Diffusion rate *D* = 0.02.

#### 2.4 The individual and population components of toxin production cost affect competition dynamics in similar ways

In the colicin system, the toxin production cost has two components: the cost, paid by all cells, of carrying the toxin producing gene (*v_p_*), which lowers the growth rate, and the cost of random cell lysis (*d_l_*) that is needed for toxin release. While the first type of cost slows down the growth and affects competition primarily at the edge of growing patches, the second type of cost increases the death rate randomly over the entire population and multiplies foci of toxin release. To test how these two types of cost affected the system, we ran the model in two ways while keeping *c*\* constant: 1. by varying the cost of carrying the toxin gene (*v_P_*), while maintaining the growth rates of the sensitive and the resistant strains unchanged; 2. by varying the lysis rate of the toxin-producing strain (*d_l_*) together with the amount of toxin released per lysis event (*p*). Increasing any of the two types of toxin production cost had similar effects: the center of the population trajectories shifted towards the corner of the sensitive strain in the ternary plots, making it more likely for the sensitive than for the resistant to win the competition (Supplementary Figure S6).

However, modifications of the second type of cost had a much stronger effect on population dynamics than the first when the overall cost (*v_P_* + *d_l_*) were considered. This is because increasing the lysis frequency and decreasing the amount of toxin released, even when keeping the value of *c*\* identical, changes the toxin distribution over the lattice and thereby profoundly affects the capacity of the producer to interfere with the sensitive strain.

#### 2.5 Varying the relative growth rates between competing strains can influence the competition outcome

Finally, we used our simulation model to test the potential consequences of differences in growth rates between strains in the range of the ones we observed in the experimental part (Figure 4A). We ran simulations with the same parameters as in Figure 7B, but with smaller or larger differences in relative growth rates between the three strains by changing the cost of resistance (*v_R_*) and of toxin production (*v_P_*). To support lasting coexistence, lower growth rate differences needed to be compensated by a decrease in the expected retention of toxin contributed by each producer cell, *c*\*; on the contrary, with higher growth rate differences between the strains, coexistence would generally require higher *c*\* (Supplementary Figure S7). This indicates that ecology-driven changes in the relative growth rates between strains, like the ones we observed between the three media in our experimental system, could lead to differences in competition outcome and population dynamics.

## Discussion

In this work, we use laboratory experiments and individual-based simulations to study how relevant ecological factors affect the competition dynamics and coexistence between toxin-producing, toxin-resistant, and sensitive *E. coli* strains. This experimental system is used as a model for cyclic dominance in the “rock-paper-scissors” evolutionary game. The main ecological factors and processes we considered are the impact of resources (i.e., amino acids and glucose) on growth and toxin production, and the accumulation, diffusion, and removal of toxin, which are known to play a major role in *E. coli’s* natural environment. The different nutrient conditions we tested by changing the CA:G ratio could emulate aspects of *E. coli’s* biphasic lifestyle inside and outside hosts [29,30] or mimic a commensal *vs* pathogenic adaptation [31,32]. In complex environments like the mammalian gut, diffusion and flow greatly vary depending on growth conditions, especially on whether cells are in a planktonic or in a sessile state within a biofilm [33,34]. Viscosity is known to promote the maintenance of high levels of toxin production in colicinogenic strains because it ensures producer cell assortment [35], but most likely also because it slows down toxin diffusion and increases the benefit the producer strain can gain from it as our model shows (Figure 6).

The three *E. coli* strains did not present obvious cyclic dominance dynamics in our bioreactors. Several factors may have contributed to this outcome. Since we only took samples once per day, the sampling frequency might have been too sparse to detect cycles with periods less than 24 hours. It is also possible that the cyclic dynamics occurring at local scales within biofilms were blurred by a relatively indiscriminate sampling method. In any case, we do not hold clear evidence that such dynamics took place in our experimental system. Although we did not frequently observe extinctions and even more rarely double extinctions, our data suggest that strain coexistence is transient in our experimental system, where trajectories often come close to boundaries. Based on our simulations, the most likely explanation for these two facts is that the toxin diffusion rate is too high to support long-term coexistence. This could be due to the thinness or lability of biofilms on the bioreactor walls, which could be incapable of retaining toxin molecules long enough in the vicinity of the producer cells. At the population level, high toxin diffusion rates could actually have a double effect: on the one hand, low toxin production rates and high toxin removal rates would prevent the producer from benefitting from the toxin and lead to the dominance of either the sensitive or the resistant like in the low CA:G medium; on the other hand, high toxin diffusion might lead to a dramatic initial reduction of the sensitive population and to the dominance of the producer (like in the high CA:G medium) or the resistant. Interestingly, in many experimental replicates, the first few days (or maybe hours) of the experiment seemed to determine the dominant strain for the following ten days. Bacteria may need to adapt to the new culture conditions in the bioreactor before starting to grow and reproduce efficiently, which could cause a drop in population size and an amplification of stochastic effects during the first hours of the run. Such effects would not have been captured by the simulations.

In our experiments, the most dramatic difference in competition outcome was due to the nutrients: while the toxin producer dominated under high CA:G, it was largely outcompeted under lower CA:G. To understand this result, it is important to observe that the *E. coli* system of cyclic dominance is fueled by two different types of competition: 1. exploitative competition, which is competition for a shared resource (e.g. nutrients) and plays a major role in the interactions between sensitive and resistant and between resistant and producer strains; 2. interference competition, which occurs when one competitor directly harms another, like when the producer secretes toxins to kill the sensitive strain. These two types of competition present very different dynamics and respond very differently to ecological factors [23,24]. The timescale of competition is typically long for exploitative competition, whose effect, in our system, requires several generations, while it is short for interference competition, since sensitive cells can be decimated within minutes once a toxin concentration threshold is reached. In the high CA:G medium, the density of all strains, including toxin producers, and the toxin production rate are both higher (Figure 4C), leading to higher overall toxin concentrations. Moreover, the relative growth rate differences between the colicin producer and the resistant strains are also smaller in this condition. The combined effect likely helped the colicin producer strain to outcompete the sensitive strain more easily in the medium with high CA:G ratio. Nevertheless, we expect that the resistant cells remaining in communities in the high CA:G medium would eventually take over if we followed the competition dynamics for a longer duration.

A feature of our work is to focus on the biological relevance of our experimental system and model design. Instead of implementing serial transfers of colonies like in the influential work of Kerr *et al*. (2002) [2], we designed and 3D-printed mini-bioreactors where we can maintain the competing community in continuous culture, keeping some degree of spatial structure in the biofilms forming on the surfaces of the bioreactor and controlling flow. This design corresponds better to natural environments, such as the bottom of river beds, the inside of water pipes, or the human/animal gut, where biofilms are under constant mechanical strain and continuous flow. Furthermore, our system avoids the artifacts due to the periodic removal of the accumulated toxin by serial transfers of colonies. This, for example, allowed us to observe the effect of the toxin remaining in the system on the competition dynamics after the producer population had disappeared (Figure 5, low CA:G, low flow). By catching the “ghost of the colicin producers”, we were able to explain the counterintuitive result that the resistant strain increased in frequency in competition with the sensitive strain for several days under low medium flow rate after the population of the producer strain had crashed.

In previous simulation models, because toxin was rarely modeled explicitly, mobility of individuals was often introduced to allow cells at some distance to interact with each other. The models have produced interesting results, such as intermediate mobility promotes coexistence [4,36], and long-distance mobility tends to cause coexistence to break down [37,38]. The way mobility was modeled was often by swapping the position of two cells or swapping the position of a cell and an empty space. It is still controversial whether those types of movement may occur in microbial biofilms, where the extracellular polymeric substances build up a matrix-like structure that limits the free movement of cells. In our model, we did not allow the cells to move, and all interactions were through the diffusion of toxin. Our finding that intermediate toxin diffusion rate promotes coexistence draws an analogy to the previous finding that intermediate mobility of individuals promotes coexistence, but is more biologically relevant and has important implications. For example, while we know that spatial structure promotes coexistence in the *E. coli* system [2,12,39], the fact that population structure without restricted toxin diffusion does not support coexistence has not been sufficiently emphasized.

It is often challenging to predict the competition dynamics and outcome between multiple interacting species in complex environments where ecological factors and processes can influence the relative competitiveness between organisms. Recently, it has been found that fairly accurate predictions of the behavior of such complex systems can be achieved by some simple rules [40]. In our work, we found that what we called toxin retention (*c*\*), which is determined by the balance between toxin release and removal, is a useful simple indicator of the propensity of long-term coexistence between the *E. coli* strains. We found that intermediate *c\** generally promotes coexistence, although extreme toxin fluxes caused by extreme toxin release and removal rates can lead to strong population fluctuations and premature collapse of coexistence. Such simple indicators, if widely applicable, can be very useful in predicting the behavior of cyclic competition dynamics in complex natural environments. We therefore encourage empirical tests in other communities where species interact in “rock-paper-scissors” games.

Evolutionary game theory, through its development in the last 50 years since the seminal work of Maynard Smith and Price [41] that opened up this field, has provided great convenience for modeling and helping us understand complex evolutionary scenarios. Classic evolutionary game models, however, often rely on simple assumptions that ignore the mechanisms underlying the strategies, and do not consider how they might be influenced by ecological factors and processes. Because theoretical studies of evolution have been more advanced than their empirical counterpart in the early days of evolutionary game theory, simple assumptions such as the “phenotypic gambit” [42] were often necessary. However, we should not forget that ecology sets up the theatre for the evolutionary play [43,44], and evolutionary game models should not ignore ecological details. Nowadays, empirical studies of ecology and evolution are much more advanced, thanks to new technological developments like the sequencing and manipulation of genetic materials, convenient analysis of biochemical molecules and their properties, as well as 3D printing that allows us to build customized devices quickly at low costs. Theoreticians also have gained access to new analytical tools and unprecedented computational power. Equipped with these new tools and accumulating knowledge in both the empirical and theoretical studies of ecology and evolution, it should be our new goal to integrate mechanisms underlying evolutionary strategies into the models for the development of evolutionary game theory in the future.

## Supporting information

Bioreactor front view

Bioreactor side view

Bioreactor top view

Raw data 1

Raw data 2

## Acknowledgement

We would like to thank Philipp Michel for creating the 3D illustration of the bioreactor, and Camille Tinguely and Valentin Voegeli for helpful discussions. The study is funded by the Swiss National Science Foundation grant PZ00P3_180142 to D.G. and grant PZ00P3_180145 to X.L.R, and the Novartis Foundation for medical-biological Research grant 19A023 to D.G.

