## Supplementary material for "Nutrients and flow shape the cyclic dominance games between *Escherichia coli* strains": Bioreactor front view

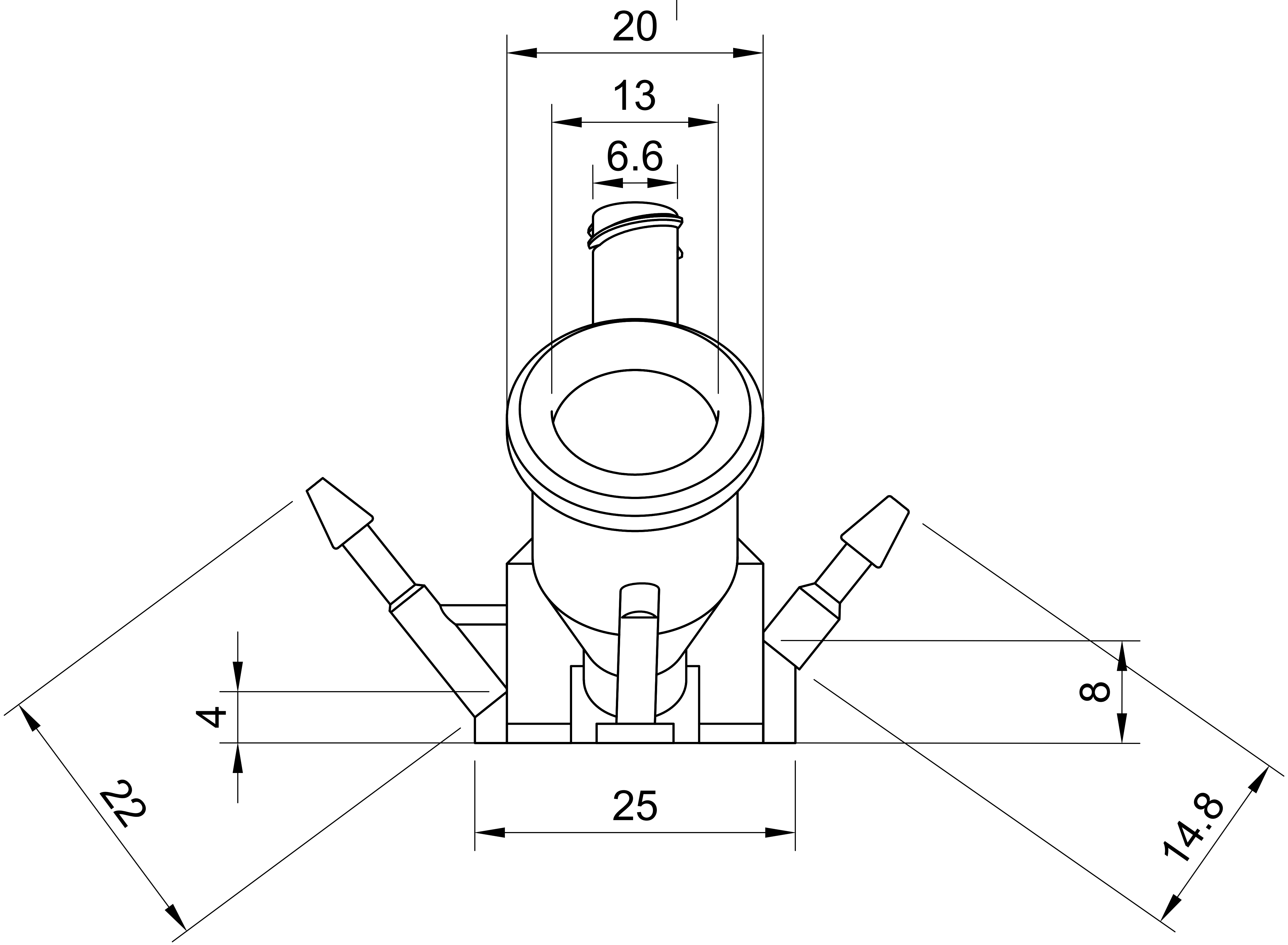


| Dept. | Technical reference | Created by | Approved by |
|---|---|---|---|
| | | Thierry Kuhn 11.08.2022 | |
| | | Document type | Document status |
| | | Title | DWG No. |
| | | Bioreactor_for_waterbath_plus_sampling_1ml_one_piece | |
| | | | Rev. / Date of issue / Sheet 1/1 |
