## Supplementary M&M for "Nutrients and flow shape the cyclic dominance games between *Escherichia coli* strains"

### Supplementary Materials and Methods

#### 1. The experiments

##### *1.1 3D-printed mini-bioreactor*

A mini-bioreactor (main text Figure 2) was designed using the Autodesk Fusion 360 software and exported as an STL file (provided in the ESM). We also provide detailed CAD drawings of the design in the ESM. It was printed with the BioMed Clear (Formlabs, RS-F2-BMCL-01) resin with a layer height of 0.1 mm using a Formlabs 3b desktop 3D printer. This resin is certified to be biocompatible, water-proof, and autoclavable. The mini-bioreactor has a working volume of 1 mL. The medium is constantly renewed through an inflow connector, connected to a medium dispenser, and an outflow connector, connected to a waste container; the flow of medium is controlled by two pumps, one for the inlet, and one for the outlet. Sterile air supply is ensured by an aeration pipe connected to a 0.22 µm mesh filter. A funnel-shaped sampling chamber capped by a rubber stopper serves for inoculation and sampling purposes.

##### *1.2 Bacterial strains*

We use three strains of the bacterium *E. coli*: the BZB1011 pColA strain that harbors the pColA plasmid and produces colicin A; the BZB1011 *btuB* strain that does not produce the toxin but is resistant to it (mutation in the colicin A receptor BtuB); and the BZB1011 strain that neither produces the toxin nor is resistant to it. The producer and resistant strains are chromosomally tagged with green and red fluorescence proteins, respectively; the cost of those markers has been shown to be negligible [1]. The three strains were cryo-preserved in 20% (v/v) glycerol at -80 °C. To reactivate the bacteria, cells were first plated on the LB agar plates (10g/L Tryptone, Oxoid; 5 g/L NaCl, Sigma-Aldrich; 5 g/L Yeast Extract, Oxoid; 15 g/L Agar, Biolife) through polygonal spreading and grown overnight at 37 °C in darkness. We then prepared bacteria suspension from the overnight culture; the three strains were inoculated into liquid LB medium and grew at 37 °C under constant rotation (300 rpm) for 16 hours. After that, the bacterial suspension was centrifuged (5 minutes at 3000 x g) and then resuspended in physiological water (9 g/L NaCl, Sigma-Aldrich). The bacterial suspension of each strain was adjusted to an optical density (at 600 nm) of 1.0.

##### *1.3 Culturing media*

We prepared three culturing media with different proportions of nutrients by supplementing the minimal *E. coli* medium M63-salts [2] with different amounts of Casamino acids (Sigma-Aldrich) and D-glucose (Carl Roth). Casamino acids contain a complex mixture of protein subunits ready for bacteria to transform into cellular building blocks. The media with high, intermediate, and low Casamino acids-to-glucose (CA:G) ratios contained 10 g/L, 3 g/L, and 1 g/L of Casamino acids and 0 g/L, 7 g/L, and 9 g/L of D-glucose, respectively. Growing in the medium with low Casamino acids requires extensive anabolism to lead to protein and lipid biosynthesis. To minimize the potentially deleterious effects of the colicin resistance mutation in poor media, we also added 1.25 mg/L of cobalamin (Fluka) to each medium.

##### *1.4 Quantification of the strain-specific growth rates under different conditions*

Cultures of the three genotypes were diluted in the three different culturing media to an optical density of 0.01. The growth of eight 200 µL replicates was monitored in a 96-well-plates incubated at 37 °C with constant agitation for 48 hours in a Biotek Synergy HT microplate reader. The optical density (600 nm) was measured once an hour. The growth rates were calculated based on data recorded during the exponential growth phase.

*1.5 Quantification of the toxin production rate under different nutrient conditions*

The 204 bps upstream of the *colA* (*cea*) gene were amplified from the pColA-CmR [3] vector by PCR using the Q5 High-Fidelity DNA polymerase (New England Biolabs) with the primers 5'-AAAGAATTCTTCCCTTCGTGATTAATAACTAAAAAC-3' and 5'-TTTAAGCTTAATCTTTCCTCTTTTTATTTTTGTTTTATGAG-3'. The fragment was cloned between the EcoRI and HindIII sites of the pPROBE-TT' [4], which encodes a promoterless green fluorescent protein (GFP) and transformed into *E. coli* DH5α-λpir by a standard heat-shock procedure. Transformants were selected on LB agar plates supplemented with tetracycline at a final concentration of 12 µg/ml. The integrity of the colA promoter sequence was verified by Sanger sequencing (Microsynth). The plasmid of interest was finally transformed into *E. coli* BZB1011 by electroporation (2.5 kV, 200 ohms, 25 µF; Bio-Rad Gene Pulser XCell). Transformants were selected on LB agar plates supplemented with tetracycline at a final concentration of 12 µg/ml and the presence of the pPROBE-TT'-P*colA* vector was verified by colony PCR.

We cultured the *E. coli* BZB1011 pProbeTT'-P*colA* in liquid LB and the three Casamino acids-glucose media with 12µg/ml tetracycline added under constant agitation (300 rpm) for 16 hours at 37 °C. We counted fluorescent and non-fluorescent cells using a Neubauer chamber (Marienfeld) under a fluorescence microscope (Leica DM4 B). We used the fluorescent intensity of producing cells in the liquid LB as a fluorescence-intensity reference. Four samples were counted for each of the three treatments. The ratio of producing cells was calculated based on total (brightfield) and fluorescent (GFP parameters) cells.

*1.6 Sensitive strain killing assay*

*E. coli* BZB1011 WT and pColA were grown in 10 ml of high CA:G medium for 24 hours and their supernatants were filtrated at 0.2 µm mesh-size filter; 100µl of each supernatant were distributed in 96-well plates in quadruplicates. *E. coli* BZB1011 WT was grown over night in high CA:G medium, washed in NaCl 0.9%, and diluted to $10^{-5}$ in NaCl 0.9%; 100 µl of cell suspension were added to the supernatants and incubated for 30 minutes before 5 µl were spotted on LB; colonies were counted the following morning.

*1.7 Experimental conditions and continuous culturing of the bacterial communities*

We implemented a factorial design of three different culturing media and two flow rates for a total of six different experimental conditions. Five replicates of the bacterial communities (each in a separate mini-bioreactor) were prepared for each experimental condition. We used two multi-channel peristaltic pumps (Longer, LP-BT100-1L) to control the inflow and outflow of the media into and out of each mini-bioreactor. The inflow rate was set to either 250 µL/h (high flow rate) or 100 µL/h (low flow rate); the outflow was kept at a constant pumping speed of 1 mL/h. Because the outlet of the mini-bioreactor was placed higher than the inlet, having a higher pumping speed at the outlet ensures sufficient air exchange and a constant volume of 1 mL in the mini-bioreactors. After connecting the mini-bioreactors, we flushed them with 100 mL fresh medium at maximum pumping speed to prevent contamination. Finally, we mixed the cell suspensions of each strain in equal proportions and pipetted 50 µl of the mixed bacterial suspensions into each mini-bioreactor by shortly removing the air filter. The mini-bioreactors were kept at 37°C in the darkness for 14 days.

##### *1.8 Quantification of the bacterial community composition*

The relative abundances of the three *E. coli* strains in each mini-bioreactor were quantified each weekday during the two weeks of experiments. A 150 µL sample was taken using a sterile syringe (1 mL, Codan) with a needle (Ø0.6 × 60 mm, Sterican) through the rubber stopper on the sampling window of each mini-bioreactor. The sample was centrifuged for 5 minutes at 2000 x g and 145 µL of the supernatant was removed. The cells were then resuspended in the rest 5 µL of the medium and placed on an agar pad. We then took pictures of the agar pad under a microscope (Leica DM4 B) in sets of three — a phase contrast picture, a picture with the GFP settings, and a picture with the RFP settings. For each sample, we took at least two pictures at different random positions on the agar pad with at least 60 cells in each picture. The pictures were then analyzed using the FIJI software (version 2.3.0, [5]) with the StarDist 2D plugin [6] to count cells of different colors. We also manually checked the results of the automatic cell detection and corrected possible detection errors in each picture.

#### 2. The simulation model

##### *2.1 Model overview*

To help better understand the experimental results, we designed a spatially explicit, individual-based simulation model. Our model was based on a $200 \times 200$ lattice with periodic boundaries, where each position contained information about its potential occupation by a bacterial cell and about the local toxin concentration (Figure 3 of the main text). We considered three bacterial cell types, corresponding to the colicin-producing, resistant, and sensitive strains of *E. coli* we used in the experiments. To initiate the simulation, we populated 1% of the positions in the lattice with a bacterium chosen at random among the three strains. The toxin concentration was set to zero everywhere. In each of the subsequent time points, we updated every position of the lattice following the order of (i) cell death and removal, (ii) toxin concentration dynamics (including the production, diffusion, and removal of colicin), and (iii) colonization of the empty positions by cell growth. The simulations were stopped either when coexistence was lost (i.e., only one cell type was left) or when they reached 300’000 steps. We implemented the model using the *Julia* programing language. The annotated simulation codes are available in the Supplementary Materials.

*2.2 Simulation of cell death*

Cells of different types can die due to different causes. Besides a baseline death probability ($d_0$) that we assumed to be identical for all cell types, the colicin-producing cells can also die from spontaneous lysis to release colicin, and the sensitive cells may be killed by colicin in the local environment. Because the resistant strain neither produces, nor can be killed by, colicin, its death probability $d_R = d_0$. The death probability of the colicin-producing strain $d_P = d_0 + d_l$, where $d_l$ denotes the spontaneous lysis probability that was set to a positive constant. In contrast to the constant death probabilities of colicin-producing and resistant cells, the death probabilities of sensitive cells are spatially and temporally heterogeneous and depend on the local colicin concentration. We denote the death probability of a sensitive cell at location $(i, j)$ as $d_{S_{i,j}} = d_0 + kc(x_{i,j}, t)$, where $k$ is a positive constant. The death probability $d_{S_{i,j}}$ is a linearly increasing function of the local colicin concentration $c$ at the position $x_{i,j}$ at time step $t$. At each time step, all bacterial cells on the lattice may die at their corresponding strain- and location-specific probabilities. The death events occurred in random order, and the dead cells were removed to leave an empty space on the lattice.

*2.3 Simulation of toxin concentration dynamics*

The lysis of every colicin-producing cell releases an equal amount of colicin into the environment. The concentration dynamics of colicin are controlled by a set of partial differential equations

$$\frac{\partial c(x_{i,j}, t)}{\partial t} = P(x_{i,j}, t) - uc(x_{i,j}, t) + D\nabla^2 c(x_{i,j}, t), \qquad \text{(Eq. 1)}$$

where $P(x_{i,j}, t)$ is the production rate of colicin, which equals a positive constant, $p$, if a colicin-producing cell was located at $x_{i,j}$ and just died at the current time step $t$; otherwise, it equals 0. The second term on the right-hand side of Eq. 1 describes the removal of colicin due to the bulk flow of culturing medium. Different values of the colicin removal rate $u$ correspond to different medium flow rates that we implemented in the experiments. We assume that the spontaneous degradation rate of colicin is negligible relative to its removal by medium flow. The diffusion rate $D$ in the last term of Eq. 1 controls the diffusion of colicin between close-by positions in the lattice. The value of $D$ could be affected by factors such as the molecular mass of different colicin molecules and the pore size distribution in the biofilm where the cells are located. Assuming that colicin diffusion occurs at much faster rates than the death and division rates of bacterial cells, we can calculate the quasi-steady-state colicin distribution at time $t$ by setting Eq. 1 to zero and solve for $c(x_{i,j}, t)$. Take the 2D finite-difference approximation of Eq. 1 and denote $c(x_{i,j}, t)$ as $c_{i,j}$ for brevity, for each position $x_{i,j}$ on the lattice, we have

$$P_{i,j} - uc_{i,j} - 4Dc_{i,j} + D(c_{i-1,j} + c_{i+1,j} + c_{i,j-1} + c_{i,j+1}) = 0, \qquad \text{(Eq. 2)}$$

where $P$ is the toxin production matrix.

We use Fourier transformations to solve the linear system of Eq. 2 efficiently, following the methods in [7]. Since we consider a $200 \times 200$ lattice with periodic boundaries, all indices in Eq. 2 should be read modulo

200, and $c_{i,j}$ is a discrete periodic function with the same period in both indices. Let $G$ be the toxin removal–diffusion kernel at position $x_{1,1}$,

$$G := \begin{pmatrix} -4D - u & D & 0 & \cdots & D \\ D & 0 & 0 & \cdots & 0 \\ 0 & 0 & 0 & \cdots & 0 \\ \vdots & \vdots & \vdots & \ddots & \vdots \\ D & 0 & 0 & \cdots & 0 \end{pmatrix}, \qquad \text{(Eq. 3)}$$

such that $G_{i,j}$ describes the diffusion and removal of toxin at each site on the lattice on the concentration of toxin at position $x_{1,1}$. The $200 \times 200$ equations defined by Eq. 2 can then be rewritten as

$$P + c^{*}G = 0, \qquad \text{(Eq. 4)}$$

where $c^{*}G$ is the circular discrete convolution of matrices $c$ and $G$. By the circular convolution theorem, $c^{*}G$ is equal to the inverse Fourier transform of the element-wise product of the individual Fourier transforms of $c$ and $G$. Therefore, we can solve the Fourier transform $F(c)$ from $F(c) \cdot F(G) = F(-P)$, and then compute the inverse Fourier transform to get the toxin concentration matrix $c$. Since the removal and diffusion rates of the molecules are constant, so is $G$. Hence, $F(G)$ needs to be calculated only once per simulation run. In each time step, we only need to calculate the Fourier transform of the toxin production matrix, dividing the result element-wise by the pre-calculated $F(G)$, and then using the inverse Fourier transform to compute the toxin concentration matrix $c$.

##### 2.4 Simulation of cell growth by colonizing empty spaces

A distinct feature of our model in contrast to typical existing lattice models of “rock-paper-scissors” games (reviewed in [8]) is the way that we modeled growth. We did not allow cells of one type to replace any other type directly; instead, the growth of cells was space-limited, corresponding to exploitative competition where space is the limiting resource, such as in typical biofilms. To grow, a cell has to have access to an empty position among the eight positions in its direct neighborhood (i.e., the Moore neighborhood [9]). Its probability of reproduction is a function of its own growth rate and the occupancy of the cells surrounding the position it attempts to divide into. Practically, for each empty position surrounded by at least one cell, the probability of this position becoming occupied by a colicin-producing, resistant, or sensitive cell was determined by the fraction of the strain’s summed growth rates over the summed growth rates of all bacterial cells in the Moore neighborhood of the empty space. The strain-specific growth rate of sensitive cells was set to $r_S = 1$. Because toxin production and resistance cost resources that can be used otherwise for growth, we assumed that the growth rate of the resistant strain $r_R = r_S - v_R$ and the growth rate of the toxin-producing strain $r_P = r_S - v_R - v_P$, where $v_R$ and $v_P$ represent the cost of colicin resistance and production, respectively. A cell cannot reproduce twice during the same timestep.

Table M1. Parameters implemented in the simulation model.

| | |
|---|---|
| $d_0$ | Baseline per time step death probability of cells |
| $d_l$ | Spontaneous lysis probability of the toxin-producing cells per time step |
| $D$ | Diffusion rate of colicin |

| | |
|---|---|
| $u$ | Removal rate of colicin by the flow of culturing medium |
| $r_S$ | Growth rate of colicin sensitive cells |
| $v_R$ | Cost on growth rate for colicin resistance |
| $v_P$ | Cost on growth rate for colicin production |
| $k$ | Killing rate of sensitive cells by colicin-induced cell lysis |
| $p$ | Amount of toxin released when a producer cell lyses |
