## Supplementary Figures for "Nutrients and flow shape the cyclic dominance games between *Escherichia coli* strains"

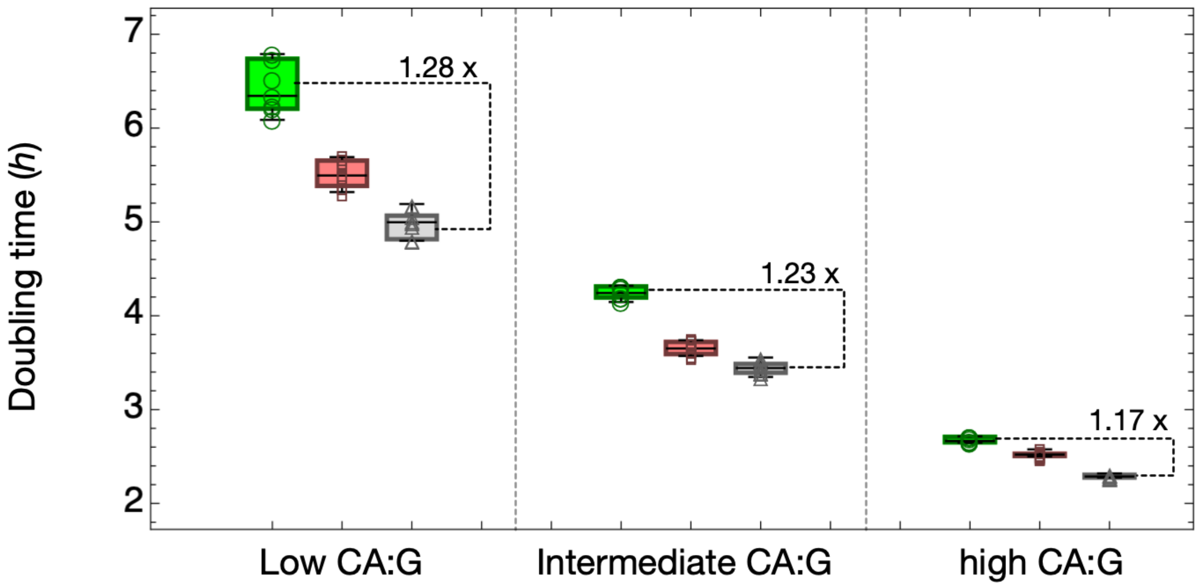


Figure S1. The doubling of three *E. coli* strains under different nutrient availability conditions. The box-whisker charts filled with green, red, and gray colors and marked with circle, square, and triangle markers represent the toxin-producer, the resistant, and the sensitive strains, respectively. The boxplot center lines show the mean, box limits show upper and lower quartiles, whiskers show the 1.5x interquartile range, and each marker corresponds to the result of an independent replicate. We have eight replicates for each bacterial strain and culturing medium combination.

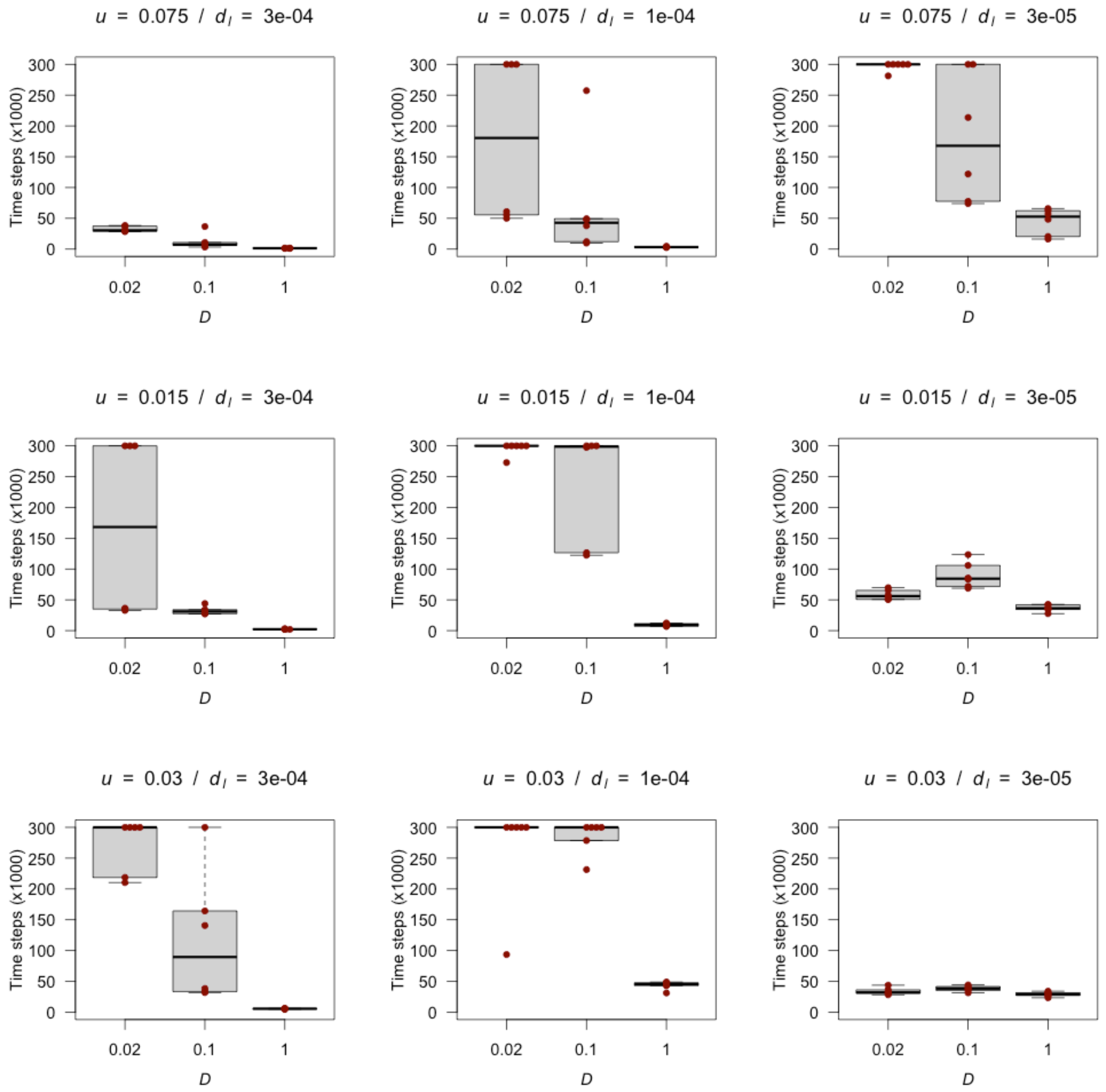


Figure S2: Coexistence times of the three strains for different toxin production and toxin removal rates with slow ($D$=0.02), fast ($D$=0.1), and instantaneous ($D$=1) toxin diffusion rates.

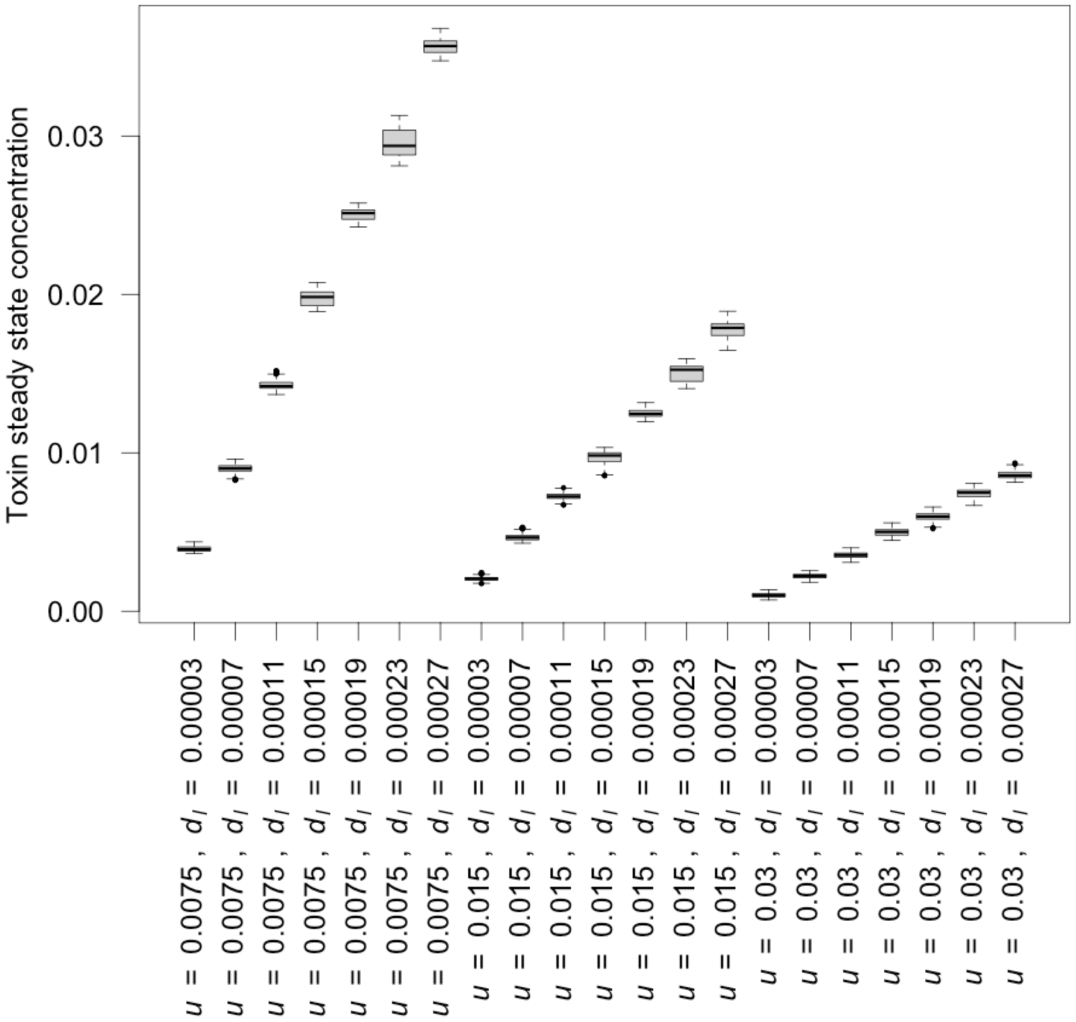


Figure S3. Steady-state toxin concentrations (between generations 700 and 1200) for different toxin production ($d_l$) and toxin removal ($u$) rates.

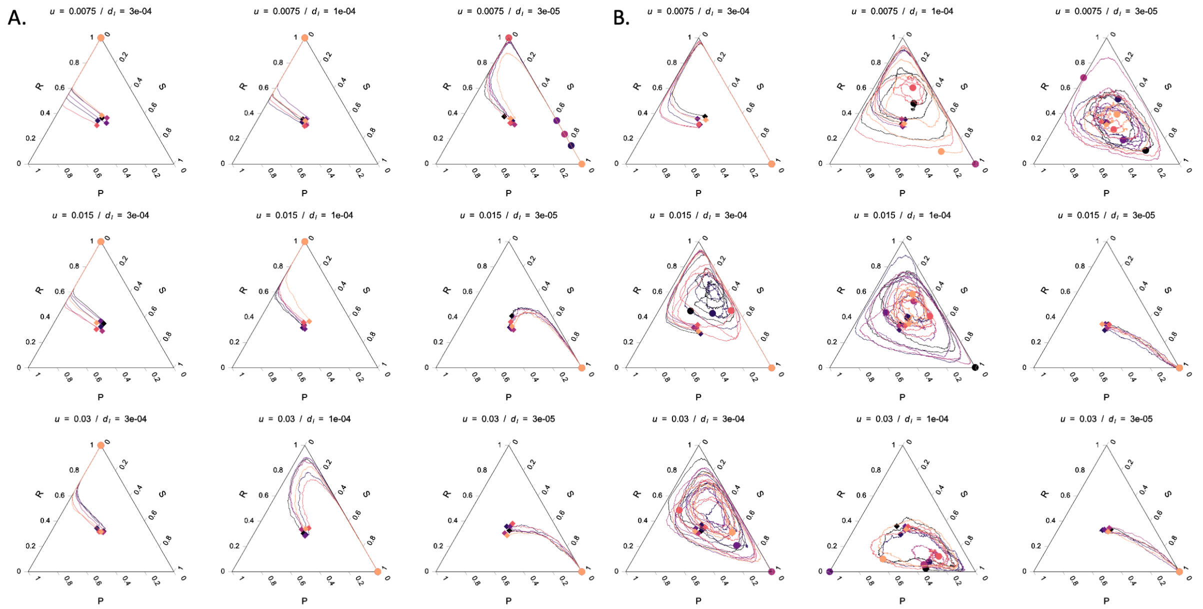


Figure S4. Population trajectory at instantaneous (panel A, $D = 1.0$) and slow diffusion (panel B, $D = 0.02$) for nine combinations of toxin production and removal rates, to be compared with the trajectories for fast diffusion shown in Figure 7B. There are six independent simulation trajectories under each condition, indicated by lines of different colors. The start point and end point of each trajectory are marked with a diamond and a circle of the same color as the trajectory, respectively. The lysis rate of toxin-producer cells, $d_l$=3e-5, 1e-4, 3e-4; toxin removal rate, $u$=0.0075, 0.015, 0.03.

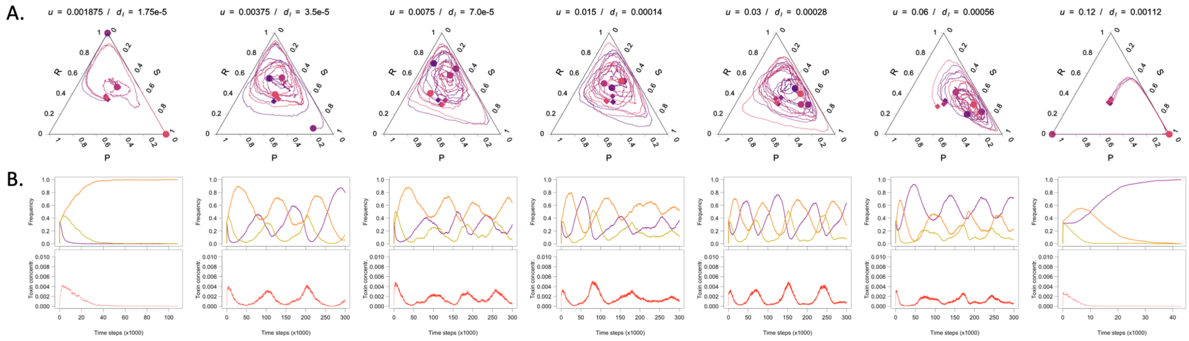


Figure S5. Impact of increasing flow and decreasing toxin production on cyclic competition patterns with toxin retention, $c^*$, kept constant. A. The values displayed above the ternary plots are the toxin removal rate ($u$) and the lysis rate of toxin-producer cells ($d_l$). B. Dynamics over time of the three strains and the toxin for one of the four replicates. Diffusion rate ($D$) is set at 0.02.

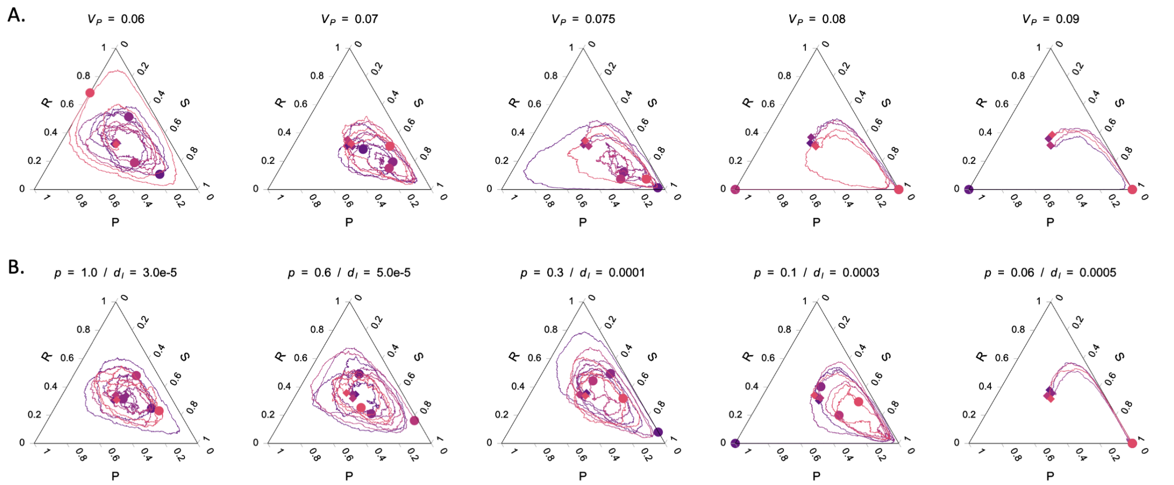


Figure S6. Impact of toxin production cost on the population trajectories. A. Trajectories for increasing toxin production costs ($v_P$), while keeping the lysis rate of producer cells ($d_l$) and toxin release per lysis event ($p$) constant. The values of $d_l$ and $p$ are set to 3e-5 and 1.0, respectively. B. Trajectories for increasing toxin-induced lysis rate ($d_l$), and correlatively adjusted toxin release per lysis event ($p$), the maximal toxin accumulation remaining the same; $v_P$ is set to 0.06. The overall toxin production cost equals $d_l + v_P$. Diffusion rate ($D$) is set at 0.1 for both panels.

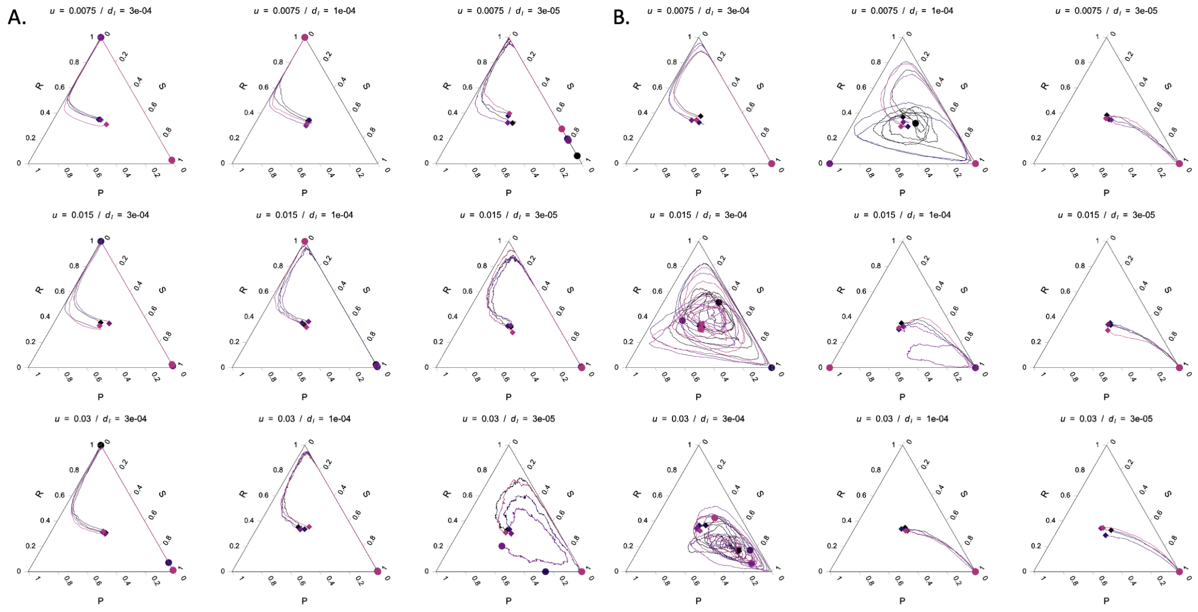


Figure S7. Population dynamics for low (A) and high (B) resistance and production costs. Simulations have been carried out with the same parameters as in Figure 7B except for $v_R = 0.01$ and $v_P = 0.02$ (panel A) or $v_R = 0.1$ and $v_P = 0.15$ (panel B). In comparison to the run presented in Figure 7B ($v_R = 0.04$ and $v_P = 0.06$), cyclic patterns tend to emerge at lower toxin concentrations in the first and at higher toxin concentrations in the second case.
